# Nurtured by nature: Considering the role of environmental and parental legacies in coral ecological performance

**DOI:** 10.1101/317453

**Authors:** Hollie M. Putnam, Raphael Ritson-Williams, Jolly Ann Cruz, Jennifer M. Davidson, Ruth D. Gates

## Abstract

The persistence of reef building corals is threatened by human-induced environmental change. Maintaining coral reefs into the future requires not only the survival of adults, but also the influx of recruits to promote genetic diversity and retain cover following adult mortality. Few studies examine the linkages among multiple life stages of corals, despite a growing knowledge of carryover effects in other systems. We provide a novel test of coral parental preconditioning to ocean acidification (OA) to better understand impacts on the processes of offspring recruitment and growth. Coral planulation was tracked for three months following adult exposure to high pCO_2_ and offspring from the second month were reciprocally exposed to ambient and high pCO_2_. Offspring of parents exposed to high pCO_2_ had greater settlement and survivorship immediately following release, retained survivorship benefits during one and six months of continued exposure, and further displayed growth benefits to at least one month post release. Enhanced performance of offspring from parents exposed to high conditions was maintained despite the survivorship in both treatments declining in continued exposure to OA. Preconditioning of the adults while they brood their larvae may provide a form of hormetic conditioning, or environmental priming that elicits stimulatory effects. Defining mechanisms of positive carryover effects, or positive trans-generational plasticity, is critical to better understanding ecological and evolutionary dynamics of corals under regimes of increasing environmental disturbance. Considering parental and environmental legacies in ecological and evolutionary projections may better account for coral reef response to the chronic stress regimes characteristic of climate change.

## Introduction

Reef-building corals are the foundation for one of the most diverse and economically important ecosystems (Bishop et al., 2011; Costanza et al., 2014). Corals are sensitive to changes in their environment that destabilize the coral-algal symbiosis and disrupt reef primary production and net ecosystem accretion. Even slight perturbations in temperature nutrients, light, salinity, and pollution, can negatively affect coral symbiosis and performance (Lesser, 2011) The steady increase in CO_2_ emissions (Pachauri et al., 2014) is heightening the magnitude and frequency of global scale environmental perturbations, exacerbating reef decline already occurring due to local pressures.

The emission of CO_2_ is driving elevated atmospheric levels, which is in turn changing the chemistry of the oceans. Ocean acidification (OA) is occurring due to the absorption of atmospheric CO_2_ by the ocean and the resulting increase and decrease in the concentrations of H^+^ and CO_3_^−2^ ion concentrations, respectively (Doney, Fabry, Feely, & Kleypas, 2009). OA energetically challenges calcifying marine organisms because they expend more energy for acid-base regulation to both maintain cellular homeostasis (Kaniewska et al., 2012; Stumpp et al., 2012) and calcify in a medium with lower CO_3_^−2^ ion concentrations. The physiological and ecological consequences of OA range from changes in intracellular pH (Gibbin, Putnam, Davy, & Gates, 2014; Gibbin, Putnam, Gates, Nitschke, & Davy, 2015), to direct reduction of calcification of coral skeletons (Erez, Reynaud, Silverman, Schneider, & Allemand, 2011), to shifts in community composition in low pH areas (Fabricius et al., 2011). Further, the interaction of multiple stressors such as temperature and OA is likely to result in synergistic and antagonistic responses in comparison to the responses driven by individual stressors (Harvey, Gwynn-Jones, & Moore, 2013; Pandolfi, Connolly, Marshall, & Cohen, 2011).

Given that local coral reef decline is now exacerbated by global issues, a greater effort is being made to better understand the sub-lethal physiological effects of stressors on critical coral reef processes such as coral reproduction and early life stages (Albright, 2011; Byrne & Przeslawski, 2013). Corals are long-lived organisms with complex reproductive life history characteristics (Baird, Guest, & Willis, 2009). Early life history research has focused heavily on the effects of anthropogenic factors (e.g., increased temperature and ocean acidification) on fertilization and development, metamorphosis, settlement, and survivorship in spawning corals (Albright, 2011; Chua, Leggat, Moya, & Baird, 2013; Chua, Leggat, Moya, & Baird, 2013; Foster, Gilmour, Chua, Falter, & McCulloch, 2015; Negri, Marshall, & Heyward, 2007; Randall & Szmant, 2009), and a variety of physiological metrics in brooding corals (Cumbo, Edmunds, Wall, & Fan, 2013; Dufault, Cumbo, Fan, & Edmunds, 2012; Putnam, Edmunds, & Fan, 2010; Rivest & Hofmann, 2014, 2015). Few studies have, however, tracked corals through multiple life history stages including larval supply, settlement and post-settlement survival and growth (Albright, 2011; Ritson-Williams, Arnold, & Paul, 2016; Ross, Ritson-Williams, Olsen, & Paul, 2013), fewer still at the cross-generational scale (Putnam & Gates, 2015), and none at the multigenerational scale. Thus the forecasts for population persistence and replenishment have largely ignored the potentially substantial plasticity driven by environmental and parental legacies (Donelson, Salinas, Munday, & Shama, 2017; Torda et al., 2017).

Phenotypic plasticity, or the generation of multiple performance outcomes within an individual as a function of environment, has the potential to act as a buffer to rapid environmental change and modulate evolutionary response (Chevin, Collins, & Lefèvre, 2013; Gibbin et al., 2017; Putnam & Gates, 2015). Multiple avenues can generate this plasticity. While intra-generational studies of phenotypic plasticity are more common, the study of trans-generational plasticity (TGP) is still in its relative infancy for marine invertebrates (Donelson et al., 2017; (Foo & Byrne, 2016); Ross, Parker, & Byrne, 2016; Torda et al., 2017). TGP occurs when environmental legacies manifest through linked life stages (i.e., the norms of reaction are shaped by the environmental conditions of prior generations). Due to the difficulty of disentangling TGP *sensu stricto* (Donelson et al., 2017; Torda et al., 2017), the plasticity exhibited between life stages can also be more generically discussed as carryover effects, which encompass the potential for TGP and developmental acclimation. The lack of data on these life stage linkages in corals is now particularly critical in light of severe ecosystem declines and the growing acknowledgement that adaptive responses are necessary for persistence under rapid climate change (Gaylord et al., 2015; Munday, Warner, Monro, Pandolfi, & Marshall, 2013; Sunday et al., 2014; Sunday, Crim, Harley, & Hart, 2011).

The ecological and evolutionary ramifications of carryover effects from parental or early life stage exposure in longer-lived, ecosystem engineering, calcifying marine invertebrates in response to OA remain to be determined. To date, early life stage exposure often appears to have negative carryover effects. Assessment of Olympia oyster performance following exposure to high pCO_2_ at the larval stage by Hettinger and coauthors revealed slower growth under high conditions. Even after the juveniles were placed in ambient grow out conditions, negative carryover effects remained at 45 days post exposure (Hettinger et al., 2012). Further assessment of the ecological performance of Olympia oyster growth in ambient field conditions four months post exposure to high pCO_2_, revealed negative carryover effects on growth rate (Hettinger et al., 2013). Some corals also appear to display negative carryover effects from stressors at the larval stage. In larvae of *Orbicella faveolata*, low salinity treatments caused decreased survival and growth after settlement (Vermeji et al., 2006). Similarly, in larvae of *Porites astreoides* early life exposure to elevated seawater temperatures caused significantly higher mortality in settled coral spat ~3 weeks post exposure in comparison to larvae raised in ambient temperatures (Ross et al., 2013). While studies in both oysters and corals identify carryover effects of early life stage exposure to stressors, this does not account for plasticity that may be induced by parental environment (i.e., TGP).

Depending on the timing of exposure, environmental conditioning can set organisms on different physiological and ecological trajectories (Foo & Byrne, 2016; Ross et al., 2016; Torda et al., 2017). Studies considering the ontogenetic connections in organisms with shorter life spans such as plankton, worms, oysters, clams, and fishes, have documented substantial, and often positive TGP (Chakravarti et al., 2016; Foo & Byrne, 2016; Gibbin et al., 2017; Lane, Campanati, Dupont, & Thiyagarajan, 2015; Miller, Watson, Donelson, McCormick, & Munday, 2012; Parker et al., 2012, 2017; Ross et al., 2016; Thor & Dupont, 2015; Zhao et al., 2018). For example, in the Sydney rock oyster, parental exposure to high pCO_2_ enhanced growth, development, and metabolism in F1 offspring exposed to the same high pCO_2_ conditions (Parker et al., 2012). Further, positive TGP was again observed in the F2 generation, in terms of development, growth, and juvenile heart rate in the trans-generational exposure line, relative to the control line (Parker, O’Connor, Raftos, Pörtner, & Ross, 2015). To date, only one study of reef building corals has demonstrated cross-generational acclimatization at the swimming planulae stage (Putnam & Gates, 2015), but the longer-term implications of this plasticity across subsequent life stages are not yet known. These responses demonstrate that understanding the potential carryover effects from parents through ontogeny (i.e., ecological ramifications of TGP) is critical to our view of organism performance on annual to decadal time scales.

The ecological implications of carryover effects have not been considered in projections of coral performance in a future of climate change, yet acclimatization is known to play a key role in intra-generational stress response (Brown & Cossins, 2011; Brown, Dunne, Goodson, & Douglas, 2000; Coles & Brown, 2003), and our initial work in corals suggests positive TGP is possible in corals (Putnam & Gates, 2015). Our current study investigates the effects of adult preconditioning on offspring ecological performance in response to OA conditions. Here, we tested the hypothesis that parental exposure of corals to high pCO_2_ during gametogenesis and/or brooding results in beneficial acclimation of the offspring. Further, we tested the hypothesis that these carryover effects are maintained beyond the larval stage, through settlement into the juvenile stage, by assessing settlement and survivorship post release and tracking survivorship and growth for six months in ambient and elevated pCO_2_ treatments.

## Materials and Methods

### Experimental Overview

The experiment consisted of four phases (Fig. 1). First, adult coral colonies were acclimated to ambient conditions in common garden tanks for 34d (Fig. 1a). Second, adults were exposed to two different pCO_2_ treatments that fluctuated with a diurnal frequency for the period that contained gametogenesis and brooding, over 3 cycles of planulation (~3 months, Fig 1b.). Third, larvae were collected from the pre-conditioned adults and exposed to the two different pCO_2_ conditions in a reciprocal fashion (Fig. 1c-e) in acrylic and mesh chambers containing a settlement tile: Survivorship and settlement of offspring during this phase were assessed after 96h. Fourth, the settled spat were tracked for survivorship and growth (Fig. 1f) in continued reciprocal exposure after one and six months post release.

**Figure 1.**
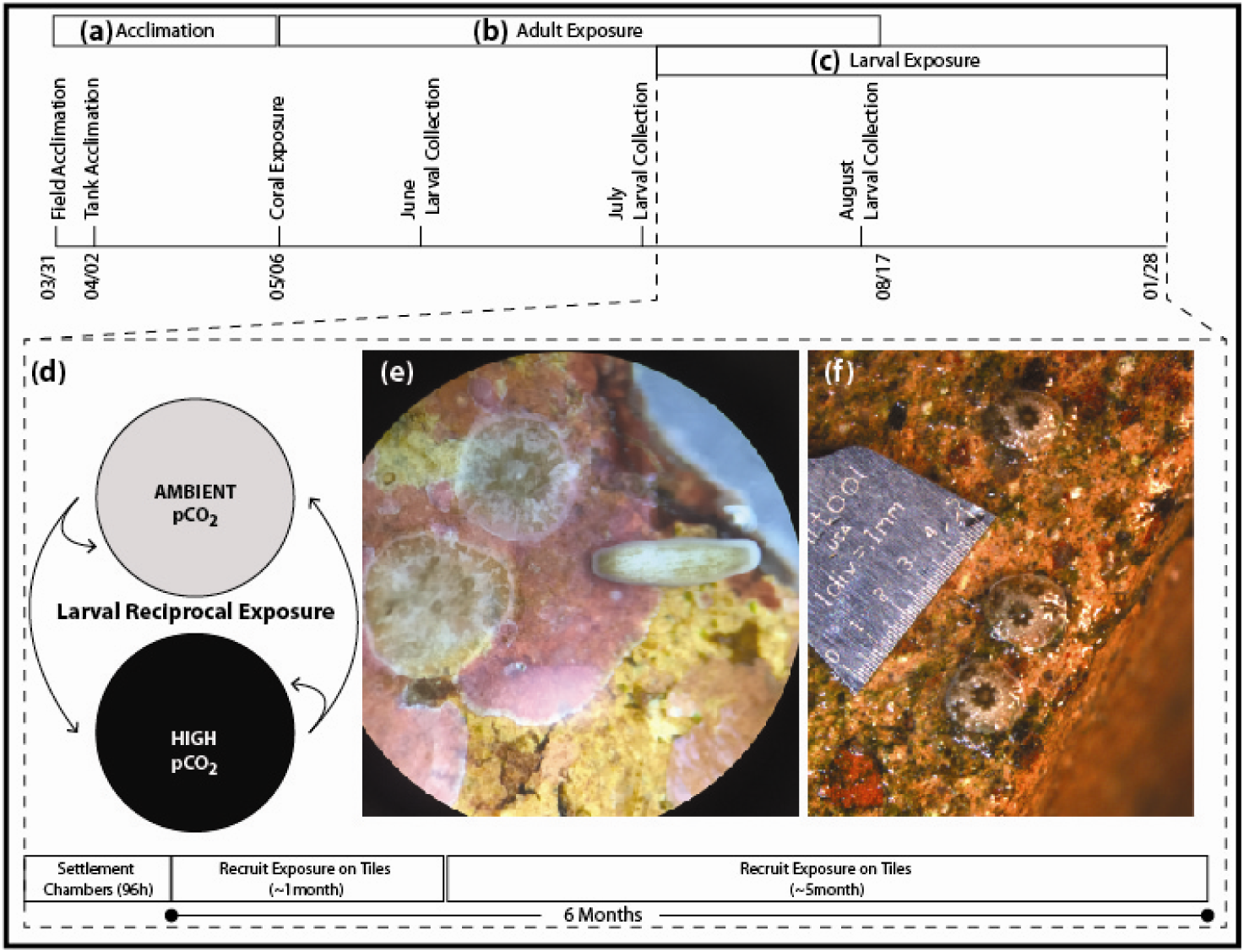
Experimental Design. (a) Coral acclimation and (b) experimental timeline to test the effects of adult preconditioning on larval release larval performance at (c, d) ambient and high pCO_2_ conditions in terms of (e) settlement, survivorship and (f) growth and survivorship were examined.

### Coral Collection and Acclimation

Adult *Pocillopora damicornis* colonies (now considered *P. acuta* based on mitochondrial open reading frame, mtORF, sequencing (Johnston, Forsman, & Toonen, 2018) were collected from the fringing reefs of southern Kāne□ohe Bay, O’ahu, Hawai□i (21.429845, −157.793604) under Special Activities Permit SAP2014 from Hawaii’s Division of Aquatic Resources (DAR). Corals were removed from the reef at the dead skeleton base to minimize direct tissue damage and were placed immediately adjacent to each other in a common garden in the field for a 3-day acclimation period (e.g., Putnam, Davidson, & Gates, 2016). Subsequently, corals (n=16) were moved to large, outdoor, sand-filtered, flow-through seawater, mesocosm tanks (~1,300L) at the Hawai□i Institute of Marine Biology, HIMB (described in Putnam et al., 2016). Irradiance in the mesocosm tanks was reduced to ~60% full irradiance using shade cloth to more closely mimic collection site *in situ* light conditions, and light readings were taken every 15 min with underwater loggers (Odyssey PAR loggers standardized to Li-Cor 192SA cosine sensor; Long et al. 2012; Fig. S1). Seawater temperature was also recorded every 15 min (Hobo Water Temp Pro v2, accuracy = 0.21°C, resolution = 0.02°C, Onset Computer Corporation, Fig. S1). Light, temperature and pH fluctuated within the tanks on natural cycles (Fig. S1). Corals acclimated to these tank conditions for 34d prior to initiation of treatments.

### Experimental Exposure

The treatment (high pCO_2_) and control conditions (ambient seawater) were maintained using a pH-stat CO_2_ injection system (Putnam et al., 2016). Ambient pH and pCO_2_ fluctuated daily (e.g., (Drupp, De Carlo, Mackenzie, Bienfang, & Sabine, 2011), driven by ambient conditions and feedbacks from photosynthesis, calcification, and respiration of the organisms on the fringing reef directly off shore of HIMB, where the seawater was obtained. The high CO_2_ treatment conditions retained natural diel variability while decreasing the pH (Fig. S1). pH probes from the pH-stat CO_2_ injection system were calibrated weekly on the NBS scale and pH and temperature were logged every 15 min in each tank throughout the duration of the experiment. The carbonate chemistry of the seawater was assessed with discrete measurements of pH (total scale), total alkalinity, temperature and salinity according to the Guide to Best Practices (Riebesell, Fabry, Hansson, Gattuso, & others, 2010). Discrete measurements were made ~daily and water samples were collected ~2x per week for each adult treatment. Given the stability of the total alkalinity and low biomass in the 1,300L tanks, sampling frequency was reduced in the 6 months of offspring exposure to ~ every 1-2 weeks, while pH, temperature, and light were tracked continuously (Fig S1). Total alkalinity samples were analyzed using open cell potentiometric titrations (Dickson, Sabine, & Christian, 2007) and assessed against certified reference materials (CRMs; A. Dickson Laboratory, UCSD; values on average <1% different from TA CRMs); all samples were corrected for any offset from the CRMs. From these measurements, the full suite of carbonate parameters was calculated with the seacarb package (v3.0.11, Gattuso et al., 2016), using the average corrected TA and salinity measured in each treatment tank (Table 1).

**Table 1.** Carbonate chemistry parameters for the different phases of the experiment (Figure 1). (a) Adult exposure (05 May 14 - 17 August 14), (b) 1 Month of offspring exposure (12 July 14 - 19 August 14), and (c) 6 Months of offspring exposure (12 July 14 - 12 January 2015). Temperature, salinity, total alkalinity, and pH were measured (N=sample size in each treatment and time point), while e remaining parameters of the carbonate system were calculated using seacarb as described in the methods.

**(a) Adult Exposures**
| Treatment | N | Temp.<br>°C | Salinity<br>psu | pH<br>Total | A<sub>T</sub><br>μmol kg<sup>-1</sup> | pCO<sub>2</sub><br>μmol kg<sup>-1</sup> | CO<sub>2</sub><br>μmol kg<sup>-1</sup> | HCO<sub>3</sub><sup>-</sup><br>μmol kg<sup>-1</sup> | CO<sub>3</sub><sup>2-</sup><br>μmol kg<sup>-1</sup> | DIC<br>μmol kg<sup>-1</sup> | Ω<sub>A</sub> |
| --- | --- | --- | --- | --- | --- | --- | --- | --- | --- | --- | --- |
| Ambient | 29 | 26.81 ± 0.12 | 33.9 ± 0.1 | 7.96 ± 0.02 | 2146 ± 5 | 482 ± 20 | 13 ± 1 | 1705 ± 14 | 178 ± 6 | 1896 ± 10 | 2.9 ± 0.1 |
| High | 28 | 26.78 ± 0.12 | 33.9 ± 0.1 | 7.71 ± 0.02 | 2151 ± 5 | 940 ± 38 | 26 ± 1 | 1878 ± 10 | 111 ± 4 | 2014 ± 7 | 1.8 ± 0.1 |

**(b) Offspring 1 Month**
| Treatment | N | Temp.<br>°C | Salinity<br>psu | pH<br>Total | A<sub>T</sub><br>μmol kg<sup>-1</sup> | pCO<sub>2</sub><br>μmol kg<sup>-1</sup> | CO<sub>2</sub><br>μmol kg<sup>-1</sup> | HCO<sub>3</sub><sup>-</sup><br>μmol kg<sup>-1</sup> | CO<sub>3</sub><sup>2-</sup><br>μmol kg<sup>-1</sup> | DIC<br>μmol kg<sup>-1</sup> | Ω<sub>A</sub> |
| --- | --- | --- | --- | --- | --- | --- | --- | --- | --- | --- | --- |
| Ambient | 8 | 27.16 ± 0.1 | 33.8 ± 0.2 | 7.9 ± 0.01 | 2138 ± 10 | 557 ± 21 | 15 ± 1 | 1747 ± 13 | 158 ± 4 | 1920 ± 12 | 2.6 ± 0.1 |
| High | 8 | 27.11 ± 0.11 | 33.8 ± 0.1 | 7.71 ± 0.03 | 2142 ± 12 | 945 ± 78 | 26 ± 2 | 1869 ± 23 | 111 ± 8 | 2005 ± 18 | 1.8 ± 0.1 |

**(c) Offspring 6 Months**
| Treatment | N | Temp.<br>°C | Salinity<br>psu | pH<br>Total | A<sub>T</sub><br>μmol kg<sup>-1</sup> | pCO<sub>2</sub><br>μmol kg<sup>-1</sup> | CO<sub>2</sub><br>μmol kg<sup>-1</sup> | HCO<sub>3</sub><sup>-</sup><br>μmol kg<sup>-1</sup> | CO<sub>3</sub><sup>2-</sup><br>μmol kg<sup>-1</sup> | DIC<br>μmol kg<sup>-1</sup> | Ω<sub>A</sub> |
| --- | --- | --- | --- | --- | --- | --- | --- | --- | --- | --- | --- |
| Ambient | 15 | 26.90 ± 0.35 | 33.8 ± 0.1 | 7.97 ± 0.04 | 2135 ± 7 | 493 ± 38 | 13 ± 1 | 1688 ± 35 | 180 ± 13 | 1882 ± 23 | 2.9 ± 0.2 |
| High | 15 | 26.92 ± 0.35 | 33.8 ± 0.1 | 7.73 ± 0.03 | 2135 ± 7 | 923 ± 70 | 25 ± 2 | 1853 ± 20 | 114 ± 8 | 1992 ± 15 | 1.8 ± 0.1 |

### Adult Exposure, Planulation, and Settlement

Eight adult corals were exposed to each treatment beginning 06 May 2014. Two colonies died very early in the high pCO_2_ treatment prior to June planulation, leaving n=6 in high pCO_2_ and n=8 in ambient pCO_2_. A common garden approach (n = 2 tanks) was chosen to maximize the similarity of experimental conditions. During the 7-8 days of larval collection each month, adult corals were removed from the tanks and separated into individual ~4.5L bowls (one colony per bowl) with flowing treatment water (i.e., the same experimental conditions as the tanks) from ~5pm to 9am to isolate the larvae released from each colony (e.g., (Putnam & Gates, 2015). The flowing seawater flushed the buoyant larvae into 800 ml tripour beakers with 150 μm mesh bottoms and the number of larvae released per colony were counted during the months of June, July and August 2014.

July 2014 was the expected seasonal peak of larval release (Fig. 2), and the larvae collected during this time were used for the offspring reciprocal exposure experiments (Fig. 1c, d). Due to the variation in timing of gametogenesis, brooding, and the sexual or asexual origin of the planulae in *Pocillopora damicornis/acuta* (Permata, Kinzie, & Hidaka, 2000; Stimson, 1978; Stoddart, 1983; Stoddart & Black, 1985), it is not possible to definitively state the exact timing of exposure of the offspring gametes or planulae within the adults. Clarifying the timing and mechanisms of carryover effects (i.e. TGP or developmental acclimation) would require, for example, exposing the adults only prior to gametogenesis, only during gametogenesis, or only during brooding. Currently the reproductive biology of *Pocillopora damicornis/acuta*, complicates this assessment. In this case, we are testing the ecological context of TGP (e.g., (Torda et al., 2017) box 2), or carryover effects.

**Figure 2.**
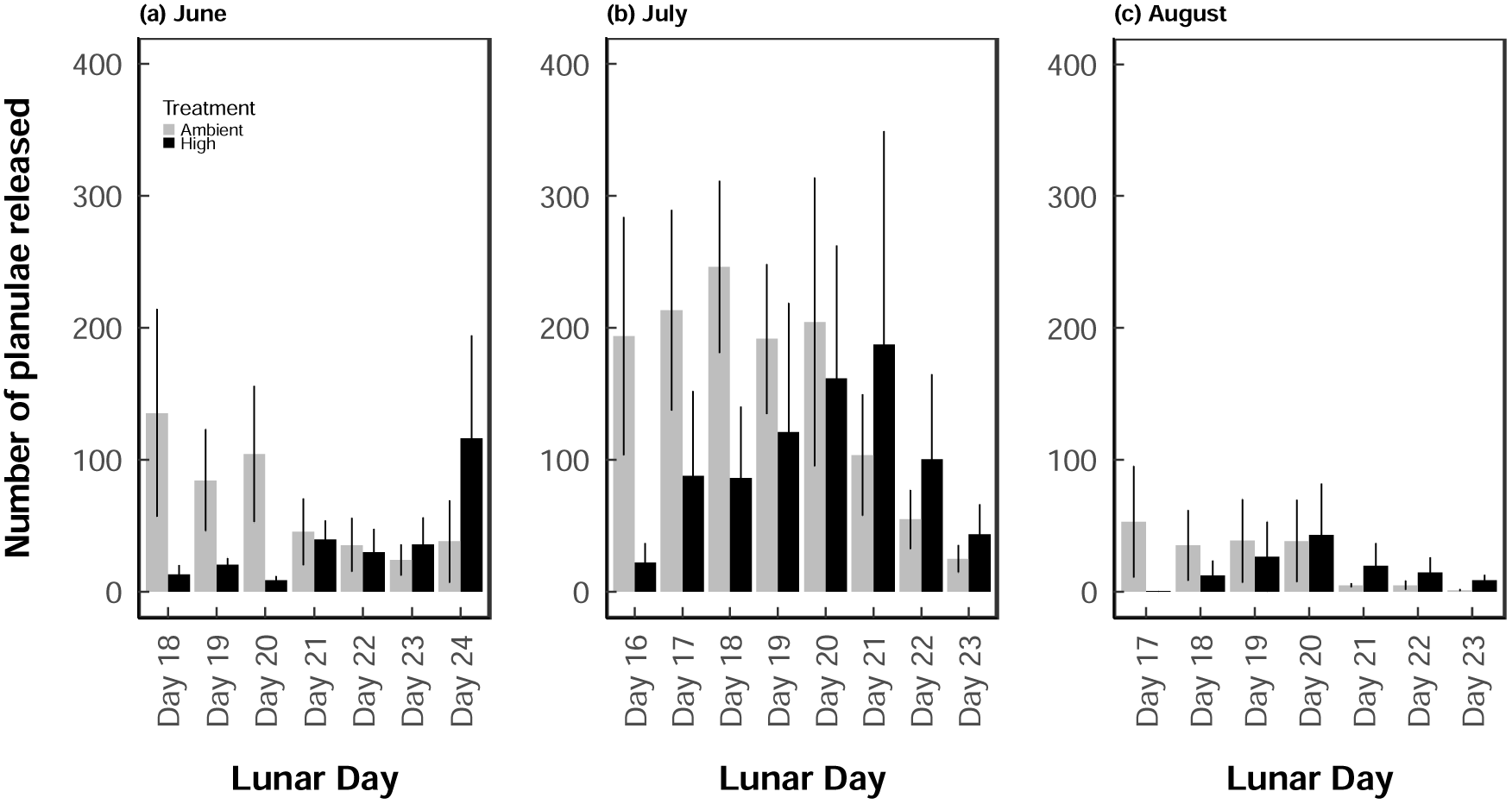
Average larval release per coral colony. Planulae release as a function of lunar day in (a) June, (b) July, and (c) August in from adult colonies exposed to ambient pCO_2_ (n=8) and adult colonies exposed to high pCO_2_ (n=6).

Groups of 16-20 larvae were tracked from each parent colony and placed in a 200ml transparent acrylic chamber with 150μm mesh at each end. To increase replication and account for variability across release date, experiments were conducted on each day of larval release from 12-18 July 2014 on newly released larvae. Each chamber contained a 4.5 × 4.5 × 1cm terracotta brick tile that had been preconditioned on the reef for 1-2 months. Conditions of pH on the fringing reef of Coconut Island in Kāne□ohe Bay can range from ~7.6 - 8.1 (Guadayol, Silbiger, Donahue, & Thomas, 2014; Silbiger, Guadayol, Thomas, & Donahue, 2014), so these tiles are not naïve to high pCO_2_. Further the major constituent was not crustose coraline algae and as such there should not be strong impacts of pCO_2_ on the tiles themselves, thereby providing relatively equal competition dynamics between the newly settled spat and tile communities. The indirect effects of pCO_2_ on substrate community, and therefore coral offspring settlement and growth, are important considerations in experimental design for interpretation of results (Albright and Langdon 2011). Chambers were placed into either ambient or high pCO_2_ treatments and settlement and survivorship were assessed after 96h. Survivorship was calculated as the total number of living larvae (both swimming and settled) in the chamber divided by the initial number of larvae added. Settlement was counted as those larvae that had settled and metamorphosed to the chamber walls, mesh, or tile divided by the initial number of larvae added to the chamber.

### Spat exposure on Tiles

After settlement and survivorship was assessed, new recruits were mapped on each tile and the tiles with settled spat were returned to the mesocosms of their settlement treatment (Figure 1, Figs. S1-S2). After both an additional month and then 6 months of total exposure, tiles were re-assessed under a dissecting scope to count survivorship (number of spat remaining alive relative to the initial amount added to the chamber). Growth of each spat was also measured by counting the number of polyps. Growth rate was calculated as: the (# of polyps - 1 primary polyp) / (# of days post settlement at month 1) for the first month’s growth rate, and as the (# of polyps at month 6 - # of polyps at month 1) / (# of days between month 1 and month 6 measurements). Spat that were fused were counted as survivors, but were not used in the growth data analysis. Settlement tile was used as the level of replication to avoid non-independence of multiple spat on a tile.

### Statistical Analysis

To test if the timing of larval release differed between the control and treatment conditions at each sampling time-point, a two-sample Kolmogorov-Smirnov test was used and release by day was treated as a continuous variable (ks.test; stats package;(R Core Team, 2016). A generalized linear mixed effects model with binomial errors (lme4 package; R Core Team, 2016) was used to test for differences in proportional settlement and survivorship between the treatment and control using a binomial distribution. Settlement data were analyzed with parental (Origin) and offspring (Secondary) treatments and their interaction as fixed effects and settlement tile as a random intercept. Survivorship and growth data were both measured at multiple time points and, thus, were modeled with the same fixed effects and interaction, but with a random intercept of settlement tile nested in time point. A model selection approach was applied and the final models were selected as those with the lowest delta AIC. Growth data were log normalized to meet model assumptions (i.e., for normal distribution) and growth data are plotted as back-transformed means and asymmetrical standard error. Full analytical details are available on GitHub (https://github.com/hputnam/HI_Pdam_Parental/releases/tag/Version_20180508) and Dryad (doi will be added upon acceptance).

## Results

### Acclimation Conditions

Corals were acclimated in tanks to fluctuating conditions (Fig. S1) similar to that on the fringing reefs of KāneLohe Bay, O’ahu, Hawai□i (Putnam et al., 2016). During the acclimation period, daily temperature ranged on average from 23.7 - 25.9°C (Fig. S1a). Between 06:00 - 19:00, Photosynthetically Active Radiation (PAR) ranged on average from 62 - 246 μmol s^−1^ m^−2^ (Fig. S1e).

### Experimental Exposure

Light and temperature varied over time within the experiment due to seasonal shifts (see supplemental Figure S1 for detailed description of tank conditions). During the adult pCO_2_ exposure, mean temperature ranged from 26.3 - 27.9°C and 26.4 - 28.0°C in ambient and treatment tanks, respectively (Fig. S1), which was on average higher than the acclimation period due to seasonal warming. pH (NBS) ranged from 7.81 - 8.06 in ambient conditions and 7.51 - 7.74 in high pCO_2_ conditions for the adult exposures. While this condition may be higher than IPCC pCO_2_ predictions for open ocean conditions (Pachauri et al., 2014), lower and more variable pH is common for coastal and reef locations (Price, Martz, Brainard, & Smith, 2012; Rivest, Comeau, & Cornwall, 2017). For example, pH conditions on the fringing reefs adjacent to Coconut Island can range from ~7.6 - 8.1 (Guadayol et al., 2014; Silbiger et al., 2014). Further modeling of the pH change in reef locations under future scenarios results in a 2.5 fold increase in reef pH variation projected with an offshore increase to 900 μatm pCO_2_ (Jury, Thomas, Atkinson, & Toonen, 2013). As such, our chosen pH conditions are ecologically relevant in terms of fluctuation and magnitude of potental pH change in the future in this dynamic reef location and do not represent extreme conditions. During the first month of larval exposure, pH ranged from 7.79 - 8.04 in ambient conditions and 7.49 - 7.74 in the high pCO_2_ treatment, and temperature, on average, ranged from 27.0 - 28.5°C and 27.0 - 28.4 °C, respectively (Fig. S1). Lastly, across the six months of juvenile exposure, pH ranged from 7.77 - 8.04 in ambient conditions and 7.52 - 7.75 in high pCO_2_ conditions and daily temperature, on average, ranged from 25.7 - 27.1°C and 25.8 - 27.0°C, respectively (Fig. S1). Discrete measurements of carbonate chemistry reveal stable TA across the 10 months and strong differences in pH and pCO_2_ between treatments (Table 1), but as these discrete measurements reflect daylight sampling only, they underestimate the treatment differences in pH and temperature clearly portrayed in continuous measurements described above (Fig. S1).

### Planulation and Settlement

Planulae release was monitored on lunar days ~16-24 for the months of June, July, and August, as planulation has been reported to occur following the full moon (lunar day ~15) for *P. damicornis* in Hawai□i (Jokiel, 1985; Richmond & Jokiel, 1984). A clear peak in planulation was observed in July, with lowest release in August (Fig. 2). There was no significant difference in the timing of planulation between treatments in either June (Fig. 2a; P > 0.05) or August (Fig. 2c; P > 0.05. The general pattern suggested a shift in timing of planulation between treatments (Fig. 2), with a delay in release more prominent in the high pCO_2_ condition in July (Fig. 2b; D = 0.625, P= 0.087).

Offspring from high CO_2_-exposed parents displayed significantly higher survivorship following 96h in the settlement chambers (P<0.016 Table S1a), supporting positive carryover effects. This increase in offspring survival from parents exposed to elevated pCO_2_ was approximately equal in both offspring treatments at time 0, with 14.5% and 15.1% greater survivorship in the ambient and high offspring treatments, respectively (Fig. 3a). The settled spat from parents preconditioned to high pCO_2_ also showed greater survivorship in both offspring treatments after one month in the reciprocal exposures (Table S1a), with preconditioned offspring having 27.1% greater survivorship at ambient offspring exposure conditions and 16.6% at high pCO_2_ exposure (Fig. 3b). After 6 months of reciprocal exposure, parental exposure to high pCO_2_ enhanced offspring survivorship by 13.6% in ambient offspring exposures, relative to spat from adults that underwent ambient preconditioning. Offspring from high pCO_2_ parents displayed reduced survivorship (7.7%) in high offspring exposure relative to spat from adults preconditioned to ambient (Fig. 3c, Table S1a). Offspring exposure to high pCO_2_ also led to lower survivorship regardless of parental preconditioning treatment (Table S1a). Averaged across parental exposures, survivorship decreased with offspring exposure to elevated pCO_2_ by 26.1% at time 0, 24.5% at month 1 and 59.9% at month 6. Survivorship declined significantly over time (P<0.001, Table S1a), with survivorship as low as 4.6% of the initial planulae by the final time point.

**Figure 3.**
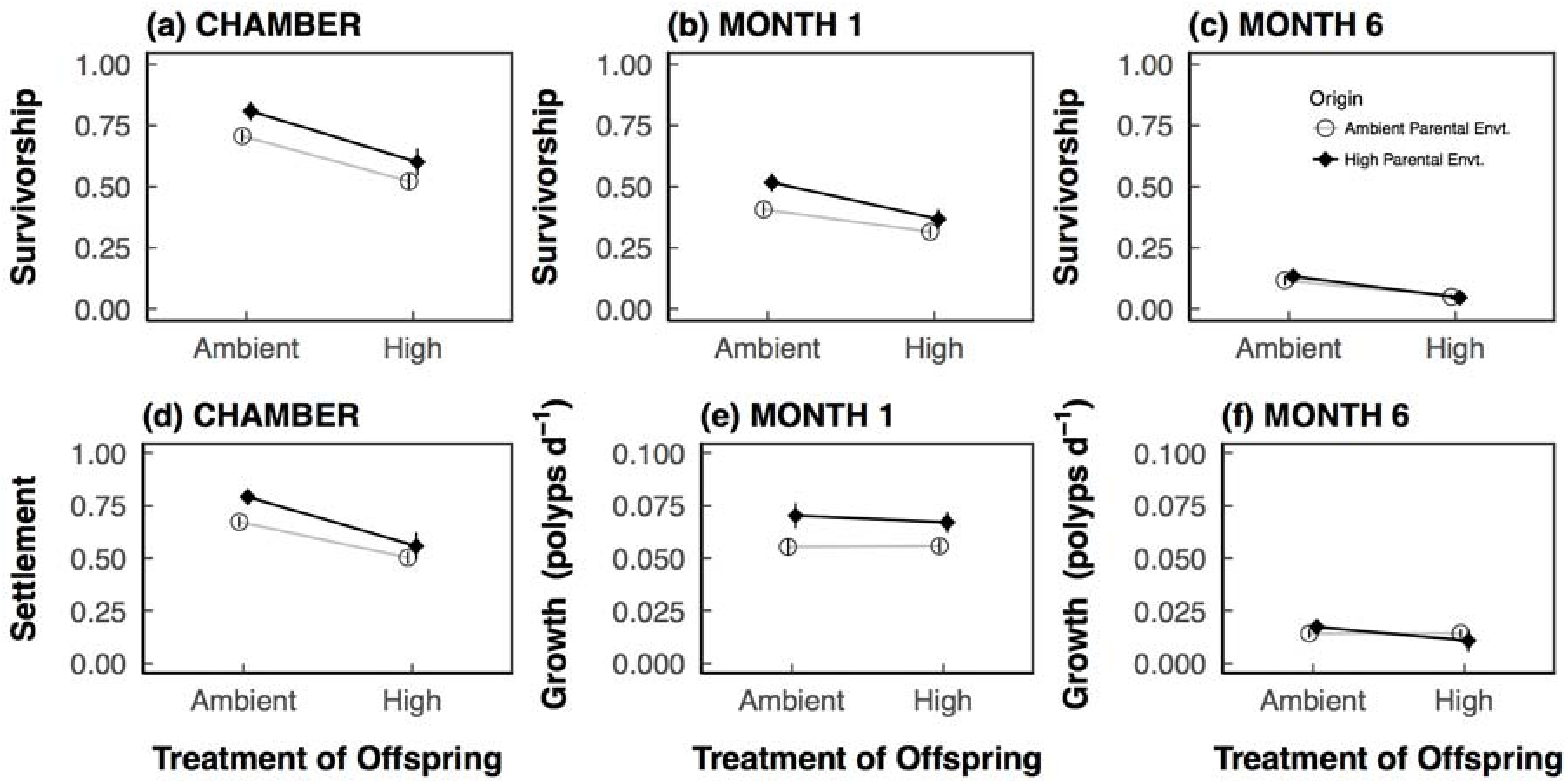
Parental preconditioning to high pCO_2_ enhances offspring performance. Reaction norm plots for (a) survivorship in the chambers, (b) survivorship at month 1, (c) survivorship at month 6, (d) settlement at time 0, (e) growth rate at month 1, and (f) growth rate at month 6. Points display data (mean±sem), while lines indicate only the direction of response. Tile sample sizes for survivorship and settlement time 0, month 1 and month 6 and growth at month 1 ranged from 7-36 (see Table S1). Growth rates were log transformed for analyses and figure displays back-transformed growth rate data.

Planulae settlement was highest when parents were exposed to elevated pCO_2_ (P=0.021; Table S1b; Fig. 3d); settlement was enhanced by 17.6% and 11% for planulae settling in ambient and high pCO_2_ offspring treatments, respectively. Despite this overall enhanced settlement of planulae from high pCO_2_ parents, mean settlement was significantly lower overall (26.1%) in the high pCO_2_ offspring treatments (P<0.001; Table S1b), regardless of parental pCO_2_ exposure (Fig. 3d).

Lastly, a trend for an Origin*Time interaction (P=0.056) was observed in spat growth, where differences in growth with parental origin were apparent at month 1 (growth was on average 1.3-fold higher in offspring from parents preconditioned to high pCO_2_; Fig. 3e), but not at month 6 (Fig. 3f). Time also had a significant effect on growth, where by the end of the experiment, at 6 months post release, growth rates were significantly lower than at month 1 (P<0.0001, Fig. 3f).

## Discussion

Here we present the first ecological assessment of trans-generational or carryover effects in reef-building corals. In our study, the ecological and fitness-related response of coral offspring was enhanced dependent on response and life-stage when their parents were exposed to a high pCO_2_ environment, with carryover effects lasting into the juvenile stage. Projections for future reef persistence are dire (Veron et al., 2009), but many do not incorporate adaptation or acclimatization (Van Hooidonk, Maynard, & Planes, 2013), likely over or under-estimating climate change effects for some stressors. When adaptation and acclimatization are considered, it is clear the trajectories differ from the worst-case scenarios (Ainsworth et al., 2016; Donner, Skirving, Little, Oppenheimer, & Hoegh-Guldberg, 2005; Logan, Dunne, Eakin, & Donner, 2014). Our work provides evidence to support the importance of the role of acclimatory processes in eco-evolutionary thinking in an era of climate change and encourage the examination of mechanisms such as hormesis and epigenetics (e.g., Costantini, 2014; Torda et al., 2017).

### Potential for tuning of reproductive timing

Our multi-life stage perspective identifies carryover effects of adult stressor exposure, with reproductive and offspring consequences. Shifts in reproductive timing can have significant consequences for offspring settlement conditions. It is possible that “bet-hedging” or environmental tuning by the parents may result in release of larvae timed to favorable conditions. For example, a shift in reproductive timing of coral planulation with change in temperature across season was recently documented (Fan et al., 2017). In this case, seasonal acclimatization in the parents resulted in change in summer planulae release timing, which minimizes planulae release during strong, likely stressful, upwelling-induced temperature fluctuations of ~10°C (Lee, Chao, Fan, Wang, & Liang, 1997). Adult tuning of larval release of offspring is also possible in our experiment. Kāne□ohe Bay is a semi-enclosed embayment that has fluctuations in physical conditions as a function of tidal cycle. Specifically, diurnal pCO_2_ fluctuations are greatest when tidal fluctuation is the lowest (Guadayol et al., 2014; Putnam, 2012; Silbiger et al., 2014) in the shallow fringing reefs of Kāne□ohe Bay. A delay in the peak of planulation from the adults preconditioned to high pCO_2_, would then correspond to the timing of lower daily tidal ranges (https://tidesandcurrents.noaa.gov/stationhome.html?id=1612480) and thus higher pCO_2_ fluctuations that are more similar to the high pCO_2_ adult conditions. Another hypothesis that could contribute to a shift in larval release under high pCO_2_ condition is adult or offspring energetic constraints. Delay in release could indicate energetic costs to maintaining adult calcification and homeostasis (Stumpp et al., 2012), potentially resulting in decreased parental investment, or increased development time necessary in offspring (Stumpp, Wren, Melzner, Thorndyke, & Dupont, 2011). Further, low pH can influence development processes such as sperm performance, fertilization success, and developmental normalcy and timing (Albright, 2011; Byrne & Przeslawski, 2013).

Given the trend for a shift in the timing of planulation during July when our offspring experiment was completed, it could be hypothesized that parental effects in the reciprocal exposure are due to slight differences in the larval cohorts by day of release (Cumbo et al., 2013; Cumbo, Fan, & Edmunds, 2012; Putnam et al., 2010; Rivest & Hofmann, 2015). For example, peaks in *Symbiodinium* density and photophysiology, and larval size, are positively correlated to peak larval release in *Pocillopora damicornis* in Taiwan (Putnam et al., 2010). These differences in physiology by day of release also translate to variation in susceptibility to changing temperature and pH in *P. damicornis* (Cumbo et al., 2013; Rivest & Hofmann, 2015). The impact of day of release in our case is likely to be minimal, given the experiment was not conducted on a single day’s larval pool, but over 7 consecutive days (Fig. 2), better representing the full range of larval phenotypes from *P. damicornis/acuta*.

### Importance of carryover effects for corals

Our work provides further evidence that parental environment matters to offspring performance in this brooding coral species, *Pocillopora acuta*. While ~1 month of exposure to increased temperature and low pH resulted in TGP in this same species (Putnam & Gates, 2015), in our current study with exposure to only OA, reaction norms were primarily parallel, with enhanced performance in those preconditioned. This may suggest that exposure to increased temperature (or the combination of temperature and OA) has more profound, or mechanistically different impacts than OA on processes involved in parental contributions or developmental acclimation (e.g., Byrne, 2012). The enhanced growth of *P. acuta* juveniles under low pH is unexpected given the commonly detrimental effect of ocean acidification on coral calcification (Kroeker, Kordas, Crim, & Singh, 2010). *Pocillopora damicornis* has, however displayed variability in sensitively of calcification in response to OA, from negative (Putnam et al., 2016) to no effects on growth (Comeau et al., 2014; Comeau, Edmunds, Spindel, & Carpenter, 2013). For instance, ocean acidification did not impact recruitment of *P. damicornis* larvae to the sides of treatment tanks in a mesocosm study also in Kāne□ohe Bay (Jokiel et al., 2008). Furthermore, physiology at the larvae stage was not strongly affected by OA (Putnam et al 2013), but was affected more so by increased temperature. Enhanced calcification in a trans-generational context has been measured in another marine invertebrate, where higher shell growth rates in offspring of the Manila clam following exposure of the parents to low pH have been observed (Zhao et al., 2018; Zhao, Schöne, Mertz-Kraus, & Yang, 2017). This positive carryover effect for growth in another marine calcifier supports our findings here with *Pocillopora acuta*, with implications for the presence and importance of TGP beyond a single coral species.

### Potential mechanisms underlying parental effects

Several hypotheses may account for the parental contribution to enhanced settlement, survivorship, and growth we documented. It is possible that adults manipulate the investment in their offspring in the form of *Symbiodinium* communities (Padilla-Gamiño, Pochon, Bird, Concepcion, & Gates, 2012), microbiome (Sharp, Distel, & Paul, 2012), size, protein, lipids, or carbohydrates (Hartmann, Marhaver, Chamberland, Sandin, & Vermeij, 2013). Further the role of mitochondrial performance has been posited as a mechanism of rapid adaptation in coral larvae (Dixon et al., 2015) and mitochondrial performance has been linked to parental environment in the trans-generational acclimatization of marine worms (Gibbin et al., 2017). These mechanisms could provide metabolic boosts or conversely detriments during this energetically demanding life stage (Edmunds, Cumbo, & Fan, 2013); e.g., the presence of clade D symbionts reduces growth in coral juveniles (Little, Van Oppen, & Willis, 2004). DNA methylation and other epigenetic mechanisms linked to gene expression regulation (Putnam et al., 2016; Putnam & Gates, 2015; Torda et al., 2017) could also provide a mechanism of heritable cross-generational priming. Identification of differential DNA methylation patterns between the offspring of exposed and unexposed oysters is an initial line of evidence for an epigenetic role in TGP for marine calcifiers (Rondon et al., 2017). The minute number of methylation changes and the lack of direct impact on RNA transcript abundance or differential splicing (Rondon et al., 2017), however, illustrates how complex these mechanisms can be. While differential provisioning of planulae symbiotically, energetically, and epigenetically is possible, the goal of examining ecological effects on the process of recruitment precluded destructive sampling for such hypotheses in our study, but they remain important considerations in future work.

Brooded embryos may experience developmental acclimation while growing within the parental polyp (Putnam & Gates, 2015). It is possible that low pH in the gastrovascular cavity (GVC) at the site of development could condition planulae for low pH when released. This hypothesis would account for the differences seen in our study between ambient and low pH preconditioned parents if the GVCs were modified differently between treatments to retain a treatment offset. To date, limited data are available for GVC pH under a variety of conditions (Agostini et al., 2011; Cai et al., 2016). The data available suggest both a high pH and low pH within the GVC and associated mesenteries (Cai et al., 2016), with no uniform picture of what planulae within the polyp are exposed to relative to external conditions. Further work with multiple coral species is necessary to disentangle the role of developmental acclimation from TGP, including experiments focused on the magnitude and timing of signals that will induce carryover effects and exposures through the F1 and F2 generation (Torda et al., 2017).

### Environmental hardening through hormetic priming

Acclimatization occurs through short-term compensatory processes including modulation of biochemical activity and gene expression in response to external stimuli, occurring on daily, seasonal, and annual scales, within a generation, and across a generation. Often implicit in the usage of the term acclimatization is the inference that acclimatory processes are beneficial and fully compensatory, *sensu* the Beneficial Acclimation Hypothesis (BAH; Huey, Berrigan, Gilchrist, & Herron, 1999), but acclimatization is not always adaptive (Edmunds, 2014; Edmunds & Gates, 2008; Huey et al., 1999). One conceptual explanation for inconsistency of BAH is the framework of hormetic priming (Costantini, 2014; Costantini, Metcalfe, & Monaghan, 2010), which deserves consideration with regards to explaining patterns of TGP. Hormesis is defined as the stimulation of function through mild exposure(s) (Southam & Ehrlich, 1943). Hormetic priming thus occurs when exposure to a sub-lethal stressor results in stimulatory response to increasing levels of a condition (e.g., Fig. 4), for example, as we have documented here in the enhanced performance of larvae of *Pocillopora acuta* following parental preconditioning. Hormesis is a biphasic response and therefore does not necessitate positive acclimation to all future exposures (Fig. 4).

**Figure 4.**
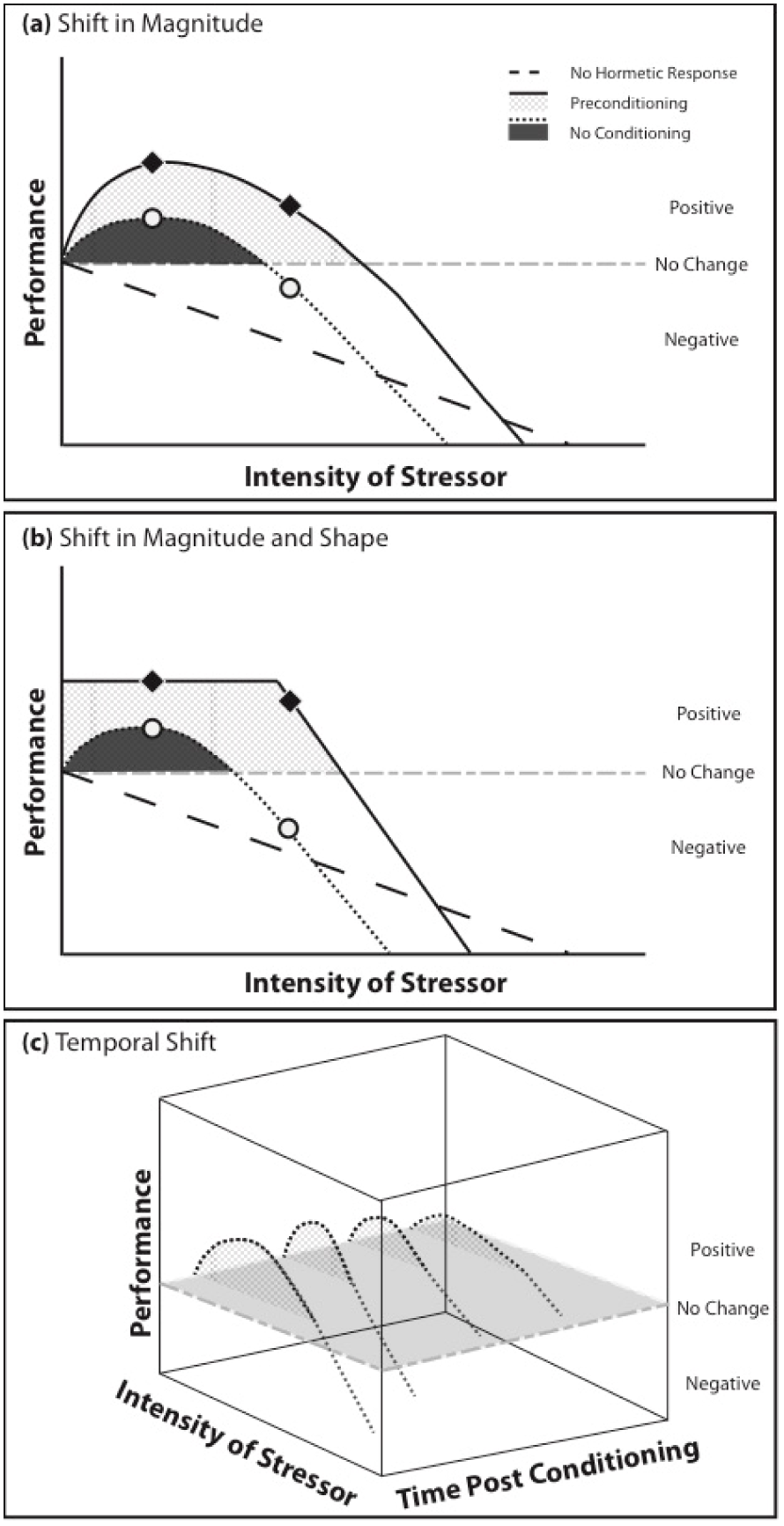
Hormetic priming may contribute to magnitude, pattern, and temporal variation in the benefits of TGP. The dashed line indicates the result of no hormetic priming, the gray dashed line is no change, the dotted line (black area) indicates a hormetic response with no conditioning, while the solid line (hashed area) indicates the hormetic response with preconditioning (Constantini et al., 2010; Lushchak 2014). Shaded areas indicate the hormetic zones, where the stimulation of performance from the increasing stressor is above the line of no change. Variation in the intensity, duration, and life stage of prior exposure may shift the A) magnitude, or B) shape of the hormetic zone. For example, in sessile benthic organisms, carryover effects from parental exposure may result in an enhanced hormetic zone relative to no conditioning. Further there may be C) temporal constraints on hormetic conditioning. For example, the hormetic zone may differ from parental exposure versus developmental exposure, or as in the case of our study, display a decline in positive carryover effects with increasing time post conditioning (as visualized by the temporal shift in hormetic zone).

Hormetic priming as a mechanism of acclimatization could explain both the beneficial and detrimental effects of increased temperature that have been documented in certain types of repeated exposures (Ainsworth et al., 2016; Brown et al., 2000; Grottoli et al., 2014), as well as variability in performance. For example, Ainsworth et al., (2016) identified a “protective trajectory” of sea surface temperatures (SST) that corals experienced, which lead to acquired thermal tolerance in *Acropora aspera*. Conversely a “repetitive bleaching trajectory”, with high frequency bleaching events that lack recovery time resulted in substantial symbiotic cell loss, bleaching, and mortality (Ainsworth et al., 2016). Here it is likely these responses are on differing sides of the zero equivalence point in terms of the biphasic nature of hormetic processes (Y axis in Fig. 4A; (Costantini et al., 2010; Lushchak, 2014). An example of hormetic priming in corals would be that of mild ROS exposure resulting in protein expression priming, thereby reducing the ROS damage in subsequent events (Brown, Downs, Dunne, & Gibb, 2002). Beyond antioxidant genes, frontloading, or the constitutive up-regulation of expression of canonical heat stress genes in *Acropora hyacinthus* samples from the high thermal variability pool in American Samoa, could be another outcome of hormetic priming, with implications for the thermal tolerance of those primed individuals (Barshis et al., 2013).

Corals in our study displayed life stage-dependent TGP. Despite the enhancement in performance seen in early stages, the parental effects of preconditioning to high pCO_2_ are absent in the offspring by 6 months post release. While maintenance of maternal effects may be expected in short lived organisms, as generation time increases there is a greater potential mismatch between parental and offspring conditions (e.g., Fig. S1). This change in parental effects over time is not unexpected due to potential maladaptive tradeoffs (Burgess & Marshall, 2014), especially the case in long-lived corals (Torda et al., 2017). Seasonal, annual, and decadal environmental changes likely elicit tradeoffs between the early life stage performance benefits of TGP and the costs of TGP the organism may incur later in life. For example, maternal exposure of soil arthropods to high temperatures resulted in increased thermal resistance at the juvenile stage, but later drove reduced F1 fecundity (Zizzari & Ellers, 2014). Further the expectation of positive TGP is linked to the predictability of the stressor (e.g., anticipatory parental effects; APE’s; Burgess & Marshall, 2014; Donelson et al., 2017). Specifically, TGP through hormetic priming would likely be optimized by high environmental autocorrelation (i.e., a strong match between parent and offspring conditions would result in extended enhancement). The greater the temporal shift from the parental environment (e.g., 6 months here Fig. S1) the less likely there is to be a benefit from parental enhancement and the more likely within generation priming would become important (e.g., Fig. 4C). Seasonal changes in physical parameters would then be expected to result in a loss of benefit from TGP as pH continues to change, in a biphasic fashion (Fig. 4C). Additionally, with respect to overall growth rates, the growth from months 2-6 would be expected to decline due to seasonal effects associated with decreases in temperature and light in the winter months (Fig. S1; Fitt, McFarland, Warner, & Chilcoat, 2000; Thornhill et al., 2011). The temporal transience of parental effects documented in our study argues that for parental effects to translate into long term “memory” more constant environmental and biological feedbacks are necessary (Ptashne, 2013).

### Implications for reef-building corals

Experiments designed to examine TGP and carryover effects provide a glimmer of hope for coral reef organisms that acclimatization may act as a buffer against a rapidly changing climate (Putnam & Gates, 2015; Torda et al., 2017). Further experiments are necessary to distinguish between true TGP, parental effects, and developmental acclimation, and their underlying mechanisms. This could be achieved in spawning coral systems, or by manipulating the timing of exposure in the brooding coral system to target isolated stages (i.e., adult, gametogenesis, brooding, and larval). Additionally, tests of the stability and later-life tradeoffs of TGP, as well as mechanisms of environmental “memory” through aspects such as mitochondrial function (Gibbin et al., 2017), DNA methylation (Putnam et al., 2016), or microbiome inheritance (Putnam, Barott, Ainsworth, & Gates, 2017; Webster & Reusch, 2017) will unveil the complex contributions of the holobiont partners to meta-organism function and acclimatization potential.

Our work challenges the paradigm of inevitable coral decline due to rapid climate change by identifying ecologically-relevant beneficial parental effects in offspring, in response to adult preconditioning to ocean acidification. We suggest hormesis, or environmental priming may conceptually explain enhanced tolerance and performance seen in acclimatization studies, as well as explaining the lack of a ubiquitous beneficial acclimation due to the biphasic nature of hormesis. With regards to a conservation and management actions (i.e. assisted evolution; van Oppen et al., 2017; van Oppen, Oliver, Putnam, & Gates, 2015), environmental priming is not a one-size-fits all phenomenon (i.e., does not always confer beneficial acclimation). Instead conditioning methods would necessitate a Goldilocks approach, as variation in cellular conditions and physiology between species requires a variety of exposures for optimal performance outcomes (Putnam et al., 2017). It is not clear, however, if the duration of the benefits would extend far beyond the exposures, as phenotype-environment mismatches increase with season and anthropogenic effects. The performance and fitness tradeoffs of acclimatization, hormetic triggers, and heritability of potential epigenetic feedbacks present promising areas of further study with respect to carryover effects and the ecological and evolutionary trajectories of reef-building corals.

## Acknowledgements

This work was supported by funding from NSF OCE-PRF 1323822 to HMP, and NSF EPS-0903833, and NSF URM 0829272. We would like to thank Rashim Khadka and the HIMB facilities staff for their assistance in this work and N. Silbiger, M. Iacchei for constructive comments on the manuscript. This is HIMB contribution number xxx and SOEST contribution number XXX.

